# Potential role of immediate early genes *Egr-1, Hr-38* and *Kakusei* in the foraging behavior and learning of in honeybees

**DOI:** 10.1101/2022.10.21.513309

**Authors:** Asem Surindro Singh, Machathoibi Takhellambam Chanu

## Abstract

The foraging behavior of honeybees is one of the most systematically composed behaviors among social insects which are admirable to watch. The main purpose of honeybee foraging is to collect food for their colony and since ancient days honeybee products have been used for various medicinal purposes (Singh and Takhellambam 2021). During foraging, honeybees gather information and transmit to their colony members regarding the location, distance, and profitability of forage sites with the help of unique movements called waggle dance. The capacities of honeybees time memory enable the foragers to return to a good food source in anticipation of the time of day. This highly intellectual, dynamic, and well-coordinated behavior of honeybees makes them to be one of the best choices of behavioral model to study various aspects of dynamic behaviors. As a result, vast knowledge in honeybee behavior has been accumulated and several recent studies immerge towards finding the underpinning regulatory biology of honeybee foraging behaviors. Immediate early genes (IE) genes are well documented neural markers and their promising roles in honeybee foraging have been demonstrated. Two of our recent studies showed three IE genes *Egr-1, Hr38* and *Kakusei* involvement during the daily foraging of honeybees. This finding has provided an avenue to further explore and identify the regulatory genes/proteins and neurons that underlie a specific behavior such as learning, memory, communication, and interaction etc. In this study we further analyze our previous published data to examine interaction of the three genes during the daily foraging of honeybees.

## Introduction

The behavior of honeybees may be considered as one of the most well-disciplined behaviors among animals. The bee colony is well maintained with different tasks that are distributed among the colony members. The queen bee is the reproductive female which specializes in egg laying and reproduces several bees of the colony, the drones are the males reserved for mating with the queen for reproduction, and the sterile workers include nursing bees that clean the hive and feed the larvae and the forgers supply food for the colony. One of the most extensively studied behaviors of honeybees is the foraging behavior. Honeybee foraging is achieved through successful shared accomplishment of several distinct behavioral features such as food search, food location, food identification, food quality checking, learning and memory, interaction, and communication among the colony members etc (Singh et al., 2018; Singh 2019; Singh et al., 2020; Singh and Takhellambam 2021; Frisch 1965; Seeley 1995). Notably the dance communication of the honeybees is considered as a symbolic way of communication in which the abstract information and concrete information are coded and decoded between individuals (Kiya et al 2012). During the dance communication bees share information with other members of the colony about the direction and distance to patches of flowers yielding nectar/pollen, water sources, new nest-site locations (Riley et al 2005; Seeley et al 2006; Frisch 1993; Dyer 2002). Such symbolic communication, like human language, requires higher brain function, which is found only in higher mammals, but not in lower animals except for honeybees (Gee 1993; Rabiah 2018). This reveals honeybees’ exceptional capability in communication skills unlike other insects or lower animals, many of their behavior features may be translated or correspond to higher animal and human behaviors.

While the importance of honeybee behavior is of great importance, the understanding of its behavior in the molecular and cellular level is still limited to few studies. Immediate early (IE) genes represent an important tool to begin searching for understanding the molecular basis of these behaviors in honeybees. Because IE genes are neural markers are also found to be neural markers in honeybee brain functions with behaviors. Any stimulus that links to membrane events and nucleus first activated IE genes and there are substantial roles of IE genes in the phenotypic changes of neurons (Loebrich and Nedivi, 2009; Hughes and Dragunow, 1995). Our three recent studies reported the possible role of three IE genes *Egr-1, Hr38* and *Kekusei* during the daily foraging honeybees and in the learning and memory during reward foraging (Singh et al., 2018; Singh et al., 2020, Singh and Takhellambam, 2021). In this study we examine the statistical examination of interaction of the three genes *Egr-1, Hr-38* and *Kakusei*, using the gene expression profile reported in the three previous studies.

## Methods

### Behavioral test and Sample collection

Behavioral test was performed inside the bee house of National Centre for Biological Sciences (NCBS), Tata Institute of Fundamental Research, Bangalore, which is an outdoor flight cage. The bees (*Apis Melifera*) were fed with pollen and 1 M sucrose solution every day from 14.00 to 17.00 hours.

Sample collection was started after the foraging bees had remembered and visited the feeders for several days. Using nectar foragers, the gene expression profiling was carried out. The detail procedures were described in our previous articles (Singh et al 2018; Singh et al 2020; Singh et al 2021).

### Sample grouping

#### During foraging

The arriving foraging bees at the feeder were caught before presenting the sucrose solution using 50 mL falcon tubes with tiny pores and immediately flash frozen in liquid nitrogen. Then, sucrose solution was presented and some of the arriving bees were gently and randomly marked on the head while drinking sucrose solution using Uni POSCA Paint Markers (Uni Mitsubishi Pencil, UK) and time count was started immediately. As the foraging bees flew repeatedly from the hive to the feeder, the marked bees were caught at different time points in 2 hours with 15 min intervals, from 14.00 hours to 16.00 hours. The time points were 15min, 30min, 45min, 60min, 75min, 90min, 105min and 120min and each time point consisted of 5 bees. The caught bees were immediately flash frozen in liquid nitrogen and stored at -80^0^C for further experiment. Figure 1 shows the partial view of the experiment.

**Figure 1.**
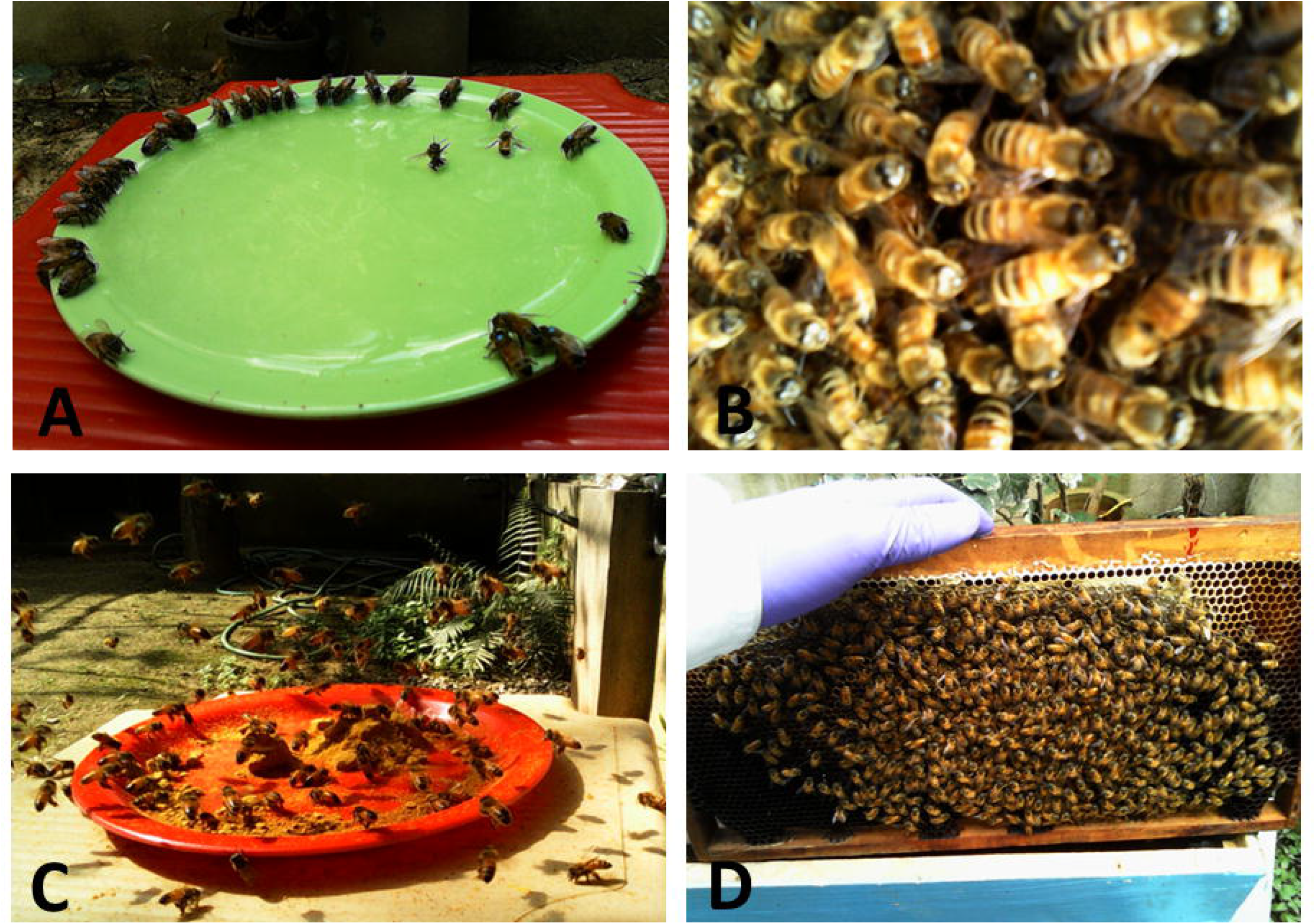
Representative picture of honeybee foraging inside the apiary. (A) honeybees feeding on 1M sucrose, (B) honeybees in the hive, (C) honeybees feeding on artificial pollen, and (D) honeybees on the honeycomb in the hive.

#### Before and after foraging

The before-foraging group consists of paint-marked foragers which were collected in the morning at 09.00 hours inside the hive, before they started foraging. In the case of after-foraging samples, the paint-marked bees were caught in the hive in the evening after the foraging of the day was over. The caught bees were immediately flash frozen in liquid nitrogen and stored at -80^0^C for further processing.

#### Un-rewarded foraging

In this group, the foraging bees were collected for 1 hour on the empty feeder plate at four time points with 15min intervals. The 0min consisted of the samples collected at 14:00 hours. After the 0min collection and the arriving bees were immediately paint-marked collected without rewarding sucrose solution at all four time points, 15min, 30min, 45min and 60min. The caught bees were immediately flash frozen in liquid nitrogen and stored at -80^0^C for further experiments.

### Gene expression profiling from the bee brains

#### Brain dissection

The frozen bees were removed from the -80°C and lyophilized at -50°C with vacuum at 0.420 mBar for 20min (Freeze Zone1 PlusTM 4.5-liter cascade Freeze Dry System, Labconco Corporation, Kanas City). The lyophilized bee head was placed in a glass chamber containing 100% ethanol placed on dry ice and the brain was dissected under the light microscope using surgical instruments. The dissected whole brain was immediately put into an Eppendorf tube on dry ice, and 500 μL Trizol (Trizol Reagent, ambion RNA, life technology) was added into it allowing it to be frozen immediately.

#### RNA and cDNA preparation

The frozen brain was allowed to thaw keeping on ice and then homogenized using electronic homogenizer (Micro-Grinder Pestle Mixer, RPI Research Products International) with pestle (Micro-Tube Sample Pestles, Research Products International). The homogenate was centrifuged at 10000g for 5min at 4^0^C, allowing total RNA, protein, DNA fractions to be formed. The upper clear fraction containing total RNA was removed gently and transferred to another tube, leaving lower DNA, tissue debris and the protein fractions undisturbed. Then cDNA was prepared from the total RNA, using the kits supplied by Invitrogen (Thermo Fisher Scientific) following manufacturer’s protocol.

#### Quantitative real time (qPCR)

qPCR was performed with the cDNA from each brain and each sample had three replicates, using 7900HT Fast Real Time PCR System (Applied Biosystem, Singapore). The total reaction volume for each sample was 10μl that contained cDNA, specific oligonucleotide primers (Sigma Aldrich) of the target genes and SYBR Green (KAPA Syber1 FAST PCR Master Mix (2X) ABI Prism1). Rp49 gene was used as endogenous control and qPCR cycles followed Applied Biosystem protocol.

### Statistical analysis

The relative gene expression level was calculated using relative standard curve method estimated by SDS 2.4 software provided with the 7900HT Fast Real system. With the help of Applied Biosystem’s ‘Guide to performing relative quantification of gene expression using real-time quantitative PCR’, standard deviation was also calculated. The fold changes at each time point were estimated relative to the level at time t0. To examine deference in gene expression level between the time points, one-way ANOVA with Turkey-Kramer post-hoc multiple comparison test was applied, and the analysis was carried out using GraphPad InStat software (Motulsky 1999). Normal distribution of each group was also examined using the D’Agostino & Pearson omnibus normality test.

To further examine interaction among the *Egr-1, Hr38* and *Kakusei* at different time points foraging of honeybees Two-way Anova was used. The statistics were carried out, analysed and interpreted with the help of GraphPad Prism 8.4.0 (www.graphpad.com) and the Handbook of Parametric and Nonparametric Statistical Procedures by David J. Sheskin (GraphPad Prism 8.4.0; www.graphpad.com). The statistics provide the percentage of the variability of gene expression among the comparing genes due to interaction between the row and column factor, the percentage due to the row factor, and the percentage due to the column factor. The remainder of the variation is among replicates (also called residual variation) not related to systematic differences between rows and columns. The null hypothesis indicates no interaction between columns (data sets) and rows whereas alternate hypothesis indicates existence of interaction. More particularly, the null hypothesis means that any systematic differences between columns are the same for each row and that any systematic differences between rows are the same for each column. Further, the null hypothesis of interaction between columns is that the mean of each column while totally ignoring the rows, is the same in the overall population, and that all differences between column means happen by chance; the column factor P value answers this question. Similarly, the null hypothesis for the interaction of the mean of each row while totally ignoring the columns, is the same in the overall population, and that all differences between row means occurred by chance; the row factor P value answers this question.

## Results

### Our recent finding on IE genes’ role in daily foraging of honeybees

Our recent studies showed that three IE genes *Egr-1, Hr38* and *Kakuei* in honeybee brain were transiently expressed and sustained for about 2 hours, during the daily food reward foraging (Singh et al., 2018; Singh and Takhelambam, 2020; Singh and Takhellambam, 2021). The upregulation declined within a few minutes when the food was not rewarded and the genes’ sustaining over expression level was due to continuous food reward. The over expression of the genes was likely due to the learned motivation by the feeder plate, but having no food rewarded the level of the genes decreased instantly. This indicates that the three genes potential involvement in foraging as well as possible link to associative learning during foraging. As the three genes showed similar patterns of expression in the same behavioral and experimental conditions, existence of interaction among them during foraging is highly possible. Using same data, here we perform the interaction analysis of the *Egr-1, Hr38* and *Kakusei* during the daily foraging of honeybees using Two-Way Anova statistics.

The significance of Interaction, Row Factor and Column Factor is at the level of P<0.0001 in each of the three experiments. This indicates that the three IE genes play a role in a collective fashion, co-operating with each other during foraging. The detail of the analysis is summarized in Table 1 and Figure 2. To check bias among the three experiments, we also performed Two Way Anova statistics for the three experiments of each gene. We observed similar level of the positive interaction (P<0.0001) indicating proximity of the three experiments. Further details of the analysis are summarized in Table 1 and Figure 3.

**Table 1.**
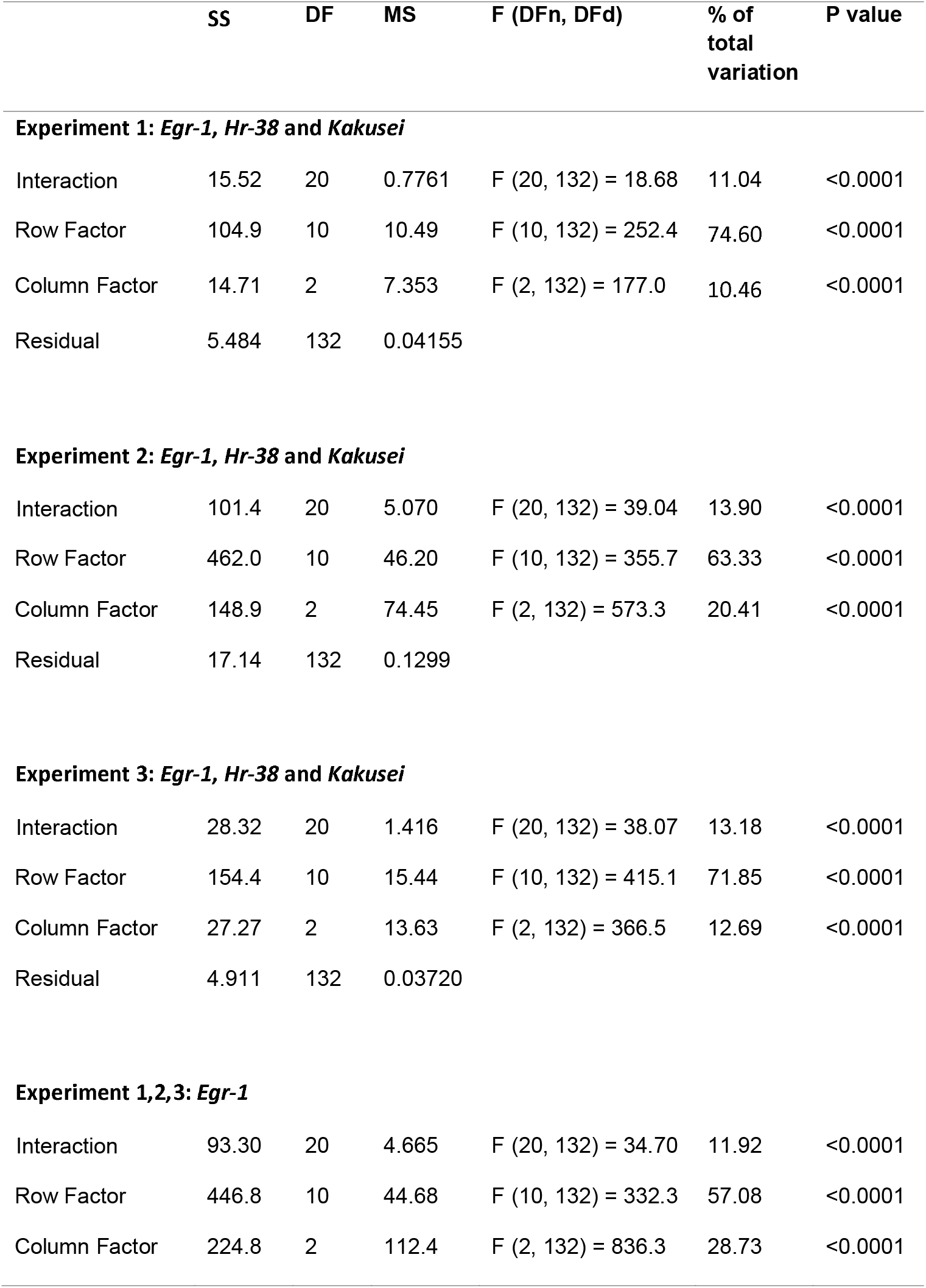
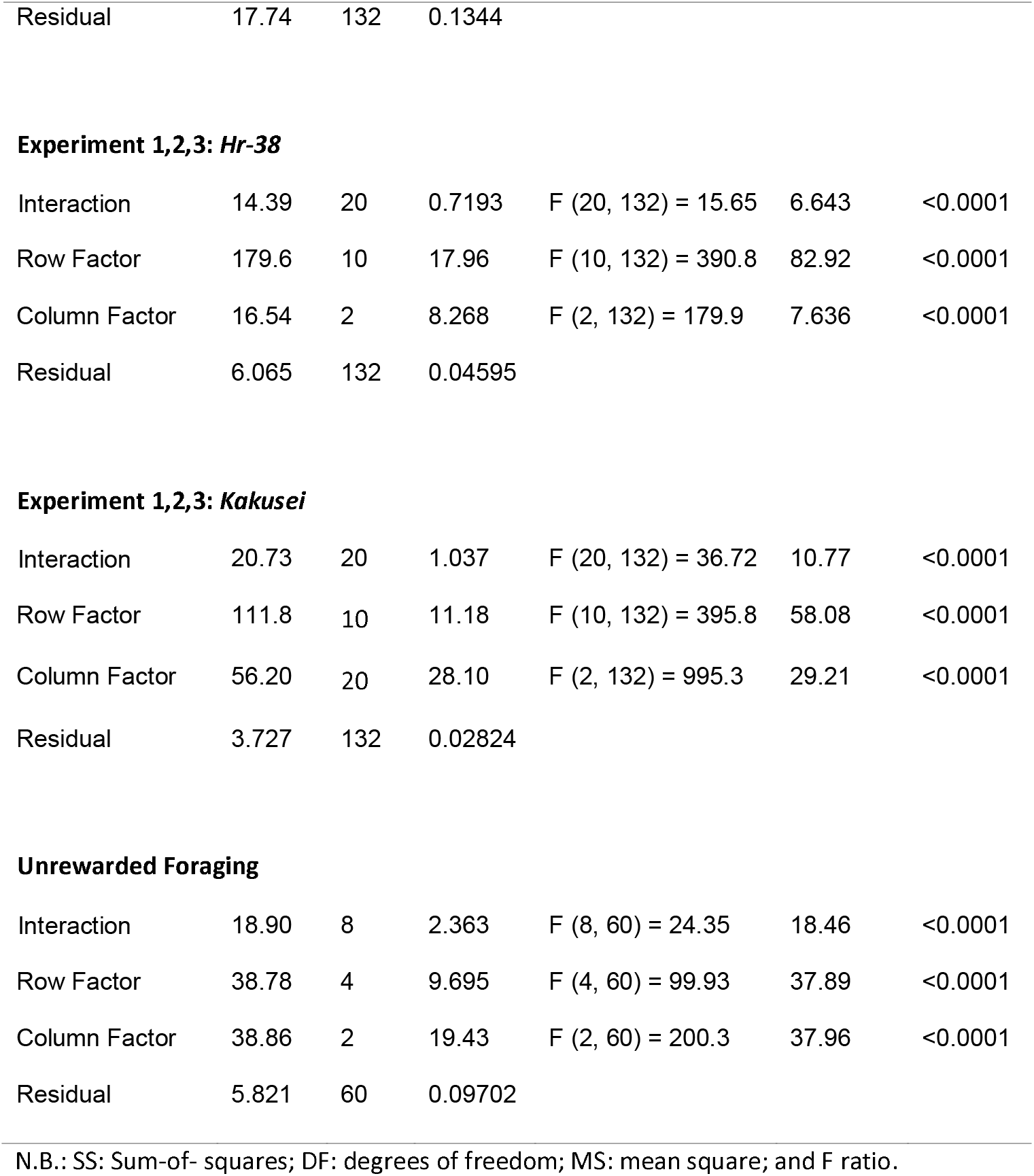
Interaction analysis of *Egr-1, Hr-38* and *Kakusei* using Two-way Anova statistics for daily rewarded and unrewarded foraging.

**Figure 2.**
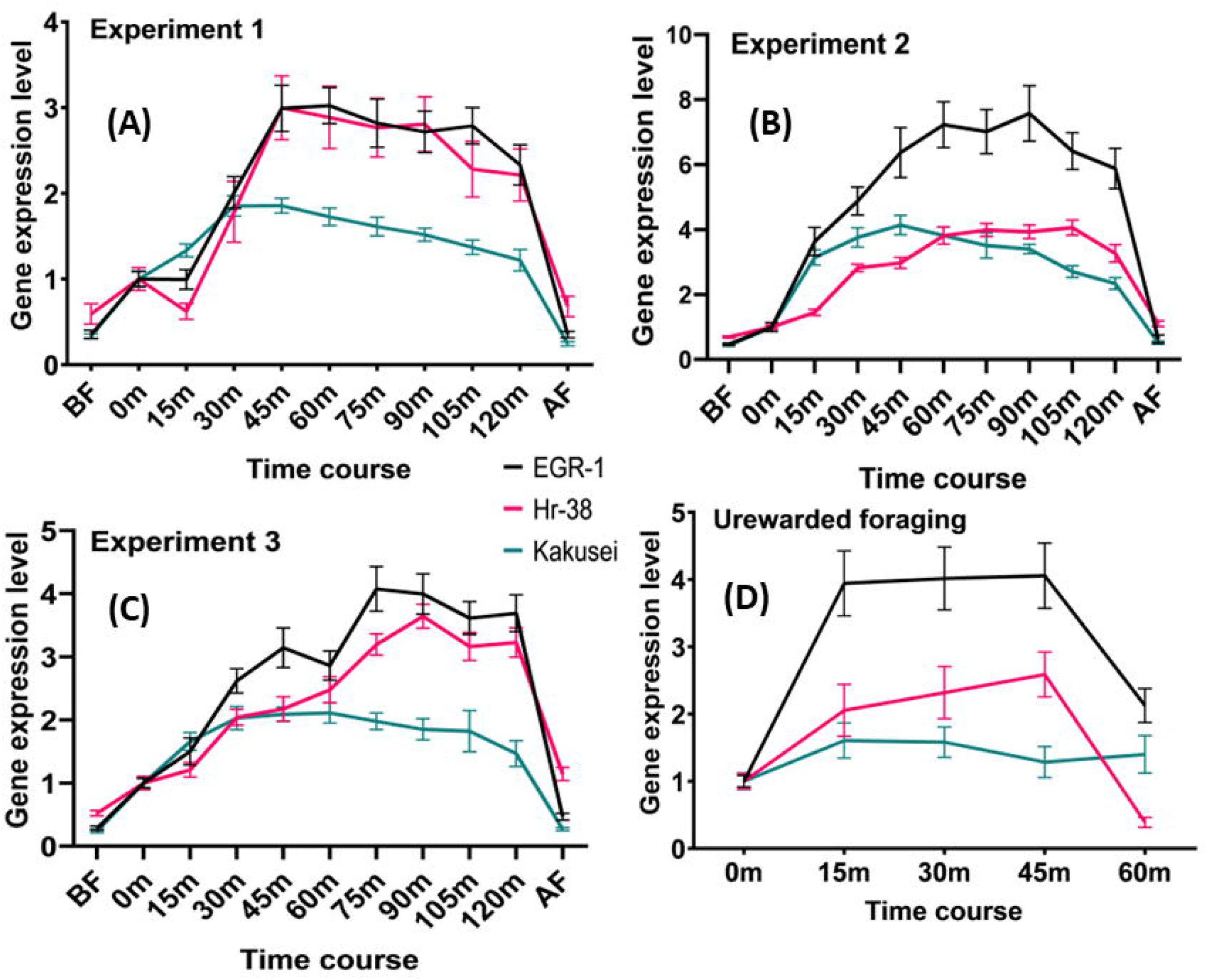
Interaction analysis for *Egr-1, Hr38* and *Kakusei*. (A) Experiment 1, (B) Experiment 2, (C) Experiment 3 and (D) Unrewarded foraging. The black line represents ***Egr-1***, red line represents *Hr38* and green line represents *Kakusei*.

**Figure 3.**
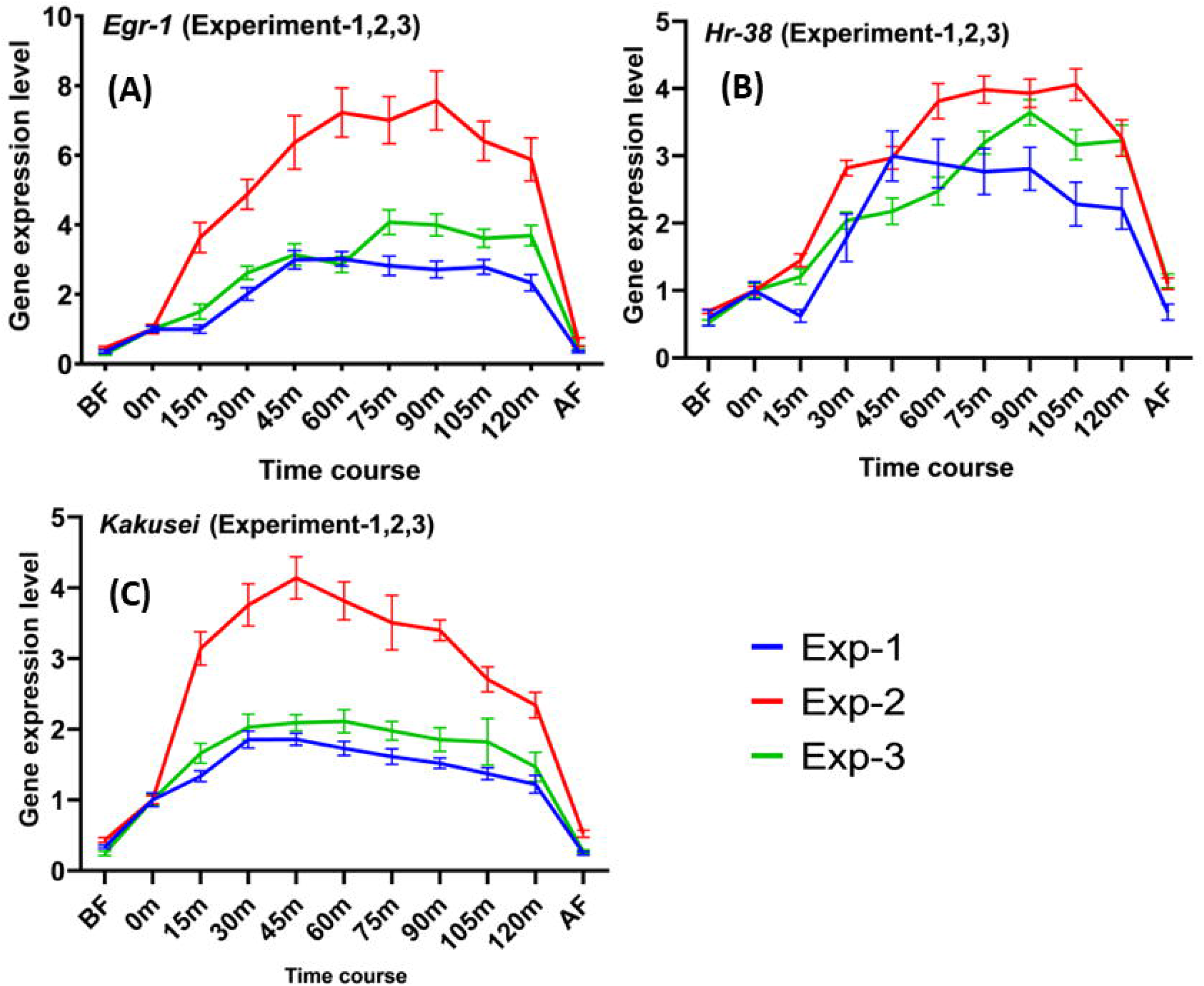
Interaction analysis for Experiment 1, Experiment 2 and Experiment 3 for *Egr-1, Hr38* and *Kakusei* respectively. (A) Experiment 1, 2 and 3 interactions for *Egr-1* (B) Experiment 2, and 3 interaction for *Hr38* (C) Experiment 1, 2 and 3 interaction for *Kakusei*. The blue line represents Experiment 1, red line represents Experiment 2 and green line represents red line represents experiment 3.

Upon the analysis of the unrewarded foraging the level of interaction among the three genes was also highly significant, as indicated by the P values of Interaction, Row Factor and Column Factor less than 0.0001. Further details of the analysis are shown in Figure 2D and Table 1.

## Discussion

Several previous works have evidenced that IE genes are neural markers, and our resent findings on *Egr-1, Hr38* and *Kakusei* on honeybee foraging have further added to it (Singh et al., 2018; Singh 2019; Singh et al., 2020; Singh and Takhellambam 2021). In our three recent findings we have shown the potential role of *Egr-1, Hr38* and *Kakusei* during the daily foraging of honeybees and associative learning during foraging. In this article we analysed whether there is any link among the three genes using the published data (Singh et al., 2018; Singh 2019; Singh et al., 2020; Singh and Takhellambam 2021). And we found a strong link among the three genes. We also tested whether there is any deviation among the repeat experiments from different colonies and we did not find any deviation but a close association among the three genes. This further strengthens the positive link among the three genes.

On the other hand, unrewarded foraging experiment further showed that the three genes are also involved in associative learning in a collective learning during foraging. It is assumable that the over expression of the genes was likely due to the learned motivation by the feeder plate, but having no food rewarded the level of the genes decreased instantly. This indicates that the three genes potential involvement in foraging as well as possible link to associative learning during foraging.

Considering the honor of novel prize to Kal Von Frisch (Von, 1993) for his incredible research and interpretation of honeybee dance, the importance of understanding of honeybee behavior has been well attributed. Over the years, one of the most extensive research areas on honeybees has been emphasized on the foraging behavior which is well worth. However, there is very little knowledge at the molecular and cellular level towards understanding the underlying mechanisms. Indeed, much can be learned from these tiny insects in terms of their well discipline specific behaviors. What fascinates us is that during foraging honeybees search for food, they identify food, taste the food, memorize food location, interact, and communicate among foragers, recruit other foragers, collect, and store food in the hive for the colony; any social animal and human would perform the similar kind of behavior. Therefore, the research findings on how honeybee foraging behaviors are regulated in the brain may be widely applicable across animal kingdoms. During foraging honeybees very well communicate each other, to be able to the underlying regulatory mechanism may be able to provide some important directions towards finding some answers towards the lack of social interaction and communication in the human neurological disorders like autism (Singh, 2014). Our finding of the positive role of Egr-1, Hr-38 and Kakusei (Singh et al., 2018; Singh 2019; Singh et al., 2020; Singh and Takhellambam 2021) and their interactions in the foraging behavior of honeybee provides an avenue towards finding the molecular pathways in the brain that connects to specific behaviors. Further research is required to give more conclusive evidence on the interactive role of *Egr-1, Hr-38* and *Kakusei* during foraging, our present data only provides statistical significance.

