## Supplementary material for "Potential role of immediate early genes *Egr-1, Hr-38* and *Kakusei* in the foraging behavior and learning of in honeybees": Table 1

**Table 1. Interaction analysis of *Egr-1, Hr-38* and *Kakusei* using Two-way Anova statistics for daily rewarded and unrewarded foraging**

| | SS | DF | MS | F (DFn, DFd) | % of total variation | P value |
|---|---|---|---|---|---|---|
| **Experiment 1: *Egr-1*, *Hr-38* and *Kakusei*** | | | | | | |
| Interaction | 15.52 | 20 | 0.7761 | F (20, 132) = 18.68 | 11.04 | <0.0001 |
| Row Factor | 104.9 | 10 | 10.49 | F (10, 132) = 252.4 | 74.60 | <0.0001 |
| Column Factor | 14.71 | 2 | 7.353 | F (2, 132) = 177.0 | 10.46 | <0.0001 |
| Residual | 5.484 | 132 | 0.04155 | | | |
| **Experiment 2: *Egr-1*, *Hr-38* and *Kakusei*** | | | | | | |
| Interaction | 101.4 | 20 | 5.070 | F (20, 132) = 39.04 | 13.90 | <0.0001 |
| Row Factor | 462.0 | 10 | 46.20 | F (10, 132) = 355.7 | 63.33 | <0.0001 |
| Column Factor | 148.9 | 2 | 74.45 | F (2, 132) = 573.3 | 20.41 | <0.0001 |
| Residual | 17.14 | 132 | 0.1299 | | | |
| **Experiment 3: *Egr-1*, *Hr-38* and *Kakusei*** | | | | | | |
| Interaction | 28.32 | 20 | 1.416 | F (20, 132) = 38.07 | 13.18 | <0.0001 |
| Row Factor | 154.4 | 10 | 15.44 | F (10, 132) = 415.1 | 71.85 | <0.0001 |
| Column Factor | 27.27 | 2 | 13.63 | F (2, 132) = 366.5 | 12.69 | <0.0001 |
| Residual | 4.911 | 132 | 0.03720 | | | |
| **Experiment 1,2,3: *Egr-1*** | | | | | | |
| Interaction | 93.30 | 20 | 4.665 | F (20, 132) = 34.70 | 11.92 | <0.0001 |
| Row Factor | 446.8 | 10 | 44.68 | F (10, 132) = 332.3 | 57.08 | <0.0001 |
| Column Factor | 224.8 | 2 | 112.4 | F (2, 132) = 836.3 | 28.73 | <0.0001 |
| Residual | 17.74 | 132 | 0.1344 | | | |
| **Experiment 1,2,3: *Hr-38*** | | | | | | |
| Interaction | 14.39 | 20 | 0.7193 | F (20, 132) = 15.65 | 6.643 | <0.0001 |
| Row Factor | 179.6 | 10 | 17.96 | F (10, 132) = 390.8 | 82.92 | <0.0001 |
| Column Factor | 16.54 | 2 | 8.268 | F (2, 132) = 179.9 | 7.636 | <0.0001 |
| Residual | 6.065 | 132 | 0.04595 | | | |
| **Experiment 1,2,3: *Kakusei*** | | | | | | |
| Interaction | 20.73 | 20 | 1.037 | F (20, 132) = 36.72 | 10.77 | <0.0001 |
| Row Factor | 111.8 | 10 | 11.18 | F (10, 132) = 395.8 | 58.08 | <0.0001 |
| Column Factor | 56.20 | 20 | 28.10 | F (2, 132) = 995.3 | 29.21 | <0.0001 |
| Residual | 3.727 | 132 | 0.02824 | | | |
| **Unrewarded Foraging** | | | | | | |
| Interaction | 18.90 | 8 | 2.363 | F (8, 60) = 24.35 | 18.46 | <0.0001 |
| Row Factor | 38.78 | 4 | 9.695 | F (4, 60) = 99.93 | 37.89 | <0.0001 |
| Column Factor | 38.86 | 2 | 19.43 | F (2, 60) = 200.3 | 37.96 | <0.0001 |
| Residual | 5.821 | 60 | 0.09702 | | | |

N.B.: SS: Sum-of- squares; DF: degrees of freedom; MS: mean square; and F ratio.
